# Leveraging single cell RNA sequencing experiments to model intra-tumor heterogeneity

**DOI:** 10.1101/427047

**Authors:** Meghan C. Ferrall-Fairbanks, Markus Ball, Eric Padron, Philipp M. Altrock

## Abstract

**PURPOSE:** Many cancers can be treated with targeted therapy. Almost inevitably, tumors develop resistance to targeted therapy, either from preexistence or by evolving new genotypes and traits. Intra-tumor heterogeneity serves as a reservoir for resistance, which often occurs due to selection of minor cellular sub-clones. On the level of gene expression, the ‘clonal’ heterogeneity can only be revealed by high-dimensional single cell methods. We propose to use a general diversity index (GDI) to quantify heterogeneity on multiple scales and relate it to disease evolution.

**METHODS:** We focused on individual patient samples probed with single cell RNA sequencing to describe heterogeneity. We developed a pipeline to analyze single cell data, via sample normalization, clustering and mathematical interpretation using a generalized diversity measure, and exemplify the utility of this platform using single cell data.

**RESULTS:** We focused on three sources of RNA sequencing data: two healthy bone marrow (BM) samples, two acute myeloid leukemia (AML) patients, each sampled before and after BM transplant (BMT), four samples of pre-sorted lineages, and six lung carcinoma patients with multi-region sampling. While healthy/normal samples scored low in diversity overall, GDI further quantified in which respect these samples differed. While a widely used Shannon diversity index sometimes reveals less differences, GDI exhibits differences in the number of potential key drivers or clonal richness. Comparing pre and post BMT AML samples did not reveal differences in heterogeneity, although they can be very different biologically.

**CONCLUSION:** GDI can quantify cellular heterogeneity changes across a wide spectrum, even when standard measures, such as the Shannon index, do not. Our approach offers wide applications to quantify heterogeneity across samples and conditions.

## INTRODUCTION

In many cancers, there still exists a critical need to understand the mechanisms of therapy resistance evolution. For example, Acute Myeloid Leukemia (AML) is an aggressive hematologic malignancy that is hallmarked by proliferation of immature myeloid cells in the bone marrow and life-threatening ineffective hematopoiesis^1^. AML is the most common adult leukemia, with an incidence of about 20,000 cases yearly and a 5-year survival of only 26%^2,3^. The diagnosis of AML requires greater than 20% of myeloid immature cells (myeloblasts) in the peripheral blood or bone marrow. The median survival of untreated AML is measured in weeks^4^. Several AML targeted therapies have been recently approved, e.g. midostaurin for FLT3 mutated patients and enasidenib for those with mutations in IDH2^5,6^. These mutations occur at rates of 25% (FLT3) and 5% (IDH2) of all AML patients and their targeted therapies are generally well tolerated relative to chemotherapeutic counterparts^7^. However, midostaurin (and even more potent FLT3 inhibitors in clinical trial^8^) does not fully eradicated the disease, leading to refractory or relapsing AML in most patients^9^. The complete response rate for enasidenib in relapse/refractory IDH2 mutated AML is less than 20%. Further refinements in patient selection are required to realize mutationally-directed therapy^5^. Little is known regarding the emerging resistance mechanism and whether targeted therapies (single or combination) against AML alone can ever be successful.

The conventional dogma postulates that therapeutic resistance occurs via the acquisition of mutations that result in clonal evolution. Emerging data suggests that these mutations are either subclonally present or present at frequencies detectable using digital PCR or ultra-deep sequencing technologies at diagnosis or prior to progression. Very low level somatic mutations are also detected in pre-leukemic states^10–13^. Somatic mutations are often present years before the diagnosis of therapy related myeloid neoplasms^14,15^. Interestingly, these mutations are commonly associated with disease progression and transformation^16^. The presence of such low-frequency genetic markers suggests that high levels of intra-tumor heterogeneity (ITH) persist over long periods of time, and that pre-existing ITH is a primary driver of future therapy resistance, while variation in transcription over time shapes the disease phenotype. A clinically relevant summary metric to describe ITH on the transcriptional level has not been developed.

scRNA-sequencing technologies can present a cost-effective method to identify transcriptomic heterogeneity and directly measure ITH. Proof of concept studies have been performed in AML, using DROP-seq that yields potentially cost-effective single cell annotations of thousands of transcripts per cell. In triple negative breast cancer, intercellular heterogeneity of gene expression programs within tumors is variable and correlates with genomic clonality^17^. A study in Chronic Myeloid Leukemia (CML) demonstrated that scRNA-seq was capable of segregating patients with discordant responses to targeted tyrosine kinase inhibitor therapy^18^. These data provide rationale to explore ITH in scRNA-seq data, and to determine whether defined measures of ITH can be predictive of progression, and eventually leveraged to mitigate progression and relapse.

Our goal is to quantify ITH in cancer such that it has maximal predictive value, in particular in hematologic malignancies. To this end we here present a platform that uses a generalized diversity index that characterizes cell population heterogeneity across a spectrum of scales (orders of diversity)^19^. These scales range from clonal richness (low order of diversity reveals the number of distinct subpopulations), to more classical measures such as Shannon or Simpson indices (intermediate order of diversity), to the number of most abundant cell types that possibly act as the key drivers of heterogeneity before transformation or perturbation by therapy (high order of diversity).

## MATERIALS AND METHODS

We created a computational and modeling approach to develop a robust statistical picture of the persistent and emerging variability in scRNA-seq data, chiefly based on drop-seq technologies; the 10x Genomics platform offered a variety of datasets that were linked to disease and treatment dynamics^20^. We specifically used the datasets of two healthy/control bone marrow mononuclear cell samples (BMMCs), two individuals with AML BMMCs sampled pre and post bone marrow transplant (BMT), to develop and test our ITH pipeline. pipelines.

First, we ran publicly available FASTQ-format files (a typical output from a drop -seq experiment) through the cellranger count pipeline and then through the cellranger aggr pipeline, to pool the samples together for comparisons during cluster analysis, interrogated through the 10x Genomics Loupe™ Cell Browser (Fig. S1). To test the robustness and valid our diversity metrics and the ITH pipeline, we extended our analysis to include additional publicly available datasets for other hematopoietic cell types (CD34+, CD14+, CD19+, and CD4+)^20^, as well as six patient normal-tumor matched lung cancer samples^21^, for which we used the same approaches and pipelines. To calculate summary metrics (outlined in **Fig. 1**), first the transcript expression data was clustered into groups of cells with similar transcript expressions (cellranger aggr). Next, we quantified the distance between each of the clusters to determine if clusters separated based on healthy or disease status (healthy vs AML). A Euclidean distance was calculated between the mean expression values for each gene, of each cluster, to establish a distance metric (**Fig. 2**).

**Figure 1.**
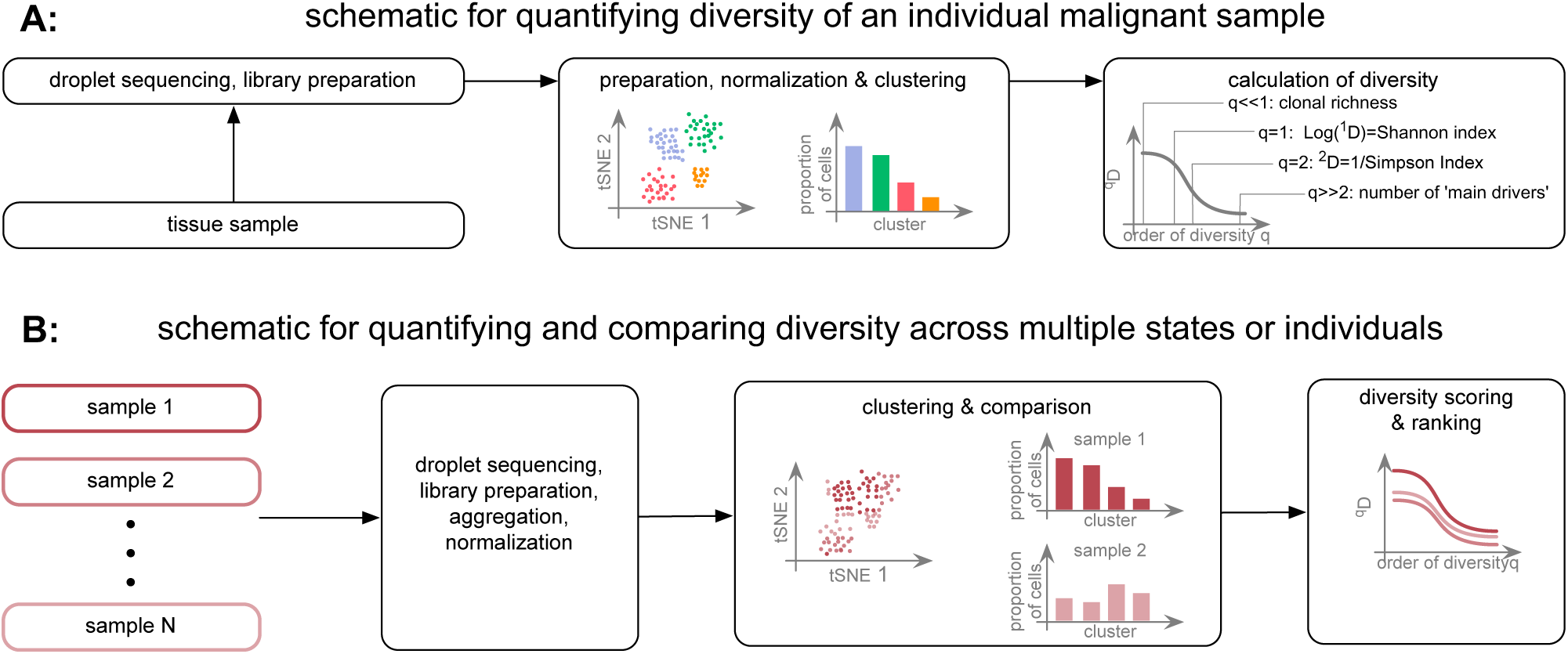
Schematic of our single cell RNA sequencing-based approach. **A:** Workflow of calculating a generalized diversity index (GDI) for a single sample. After sequencing and library preparation, normalization to reduce the number of false negatives or false positives is applied, for example using 10x-genomics platform, or clustering can then be applied (Loupe™ Cell browser, or other platforms, see Supplement), from which diversity can be calculated. **B**: A similar approach can be used when multiple samples are compared. Data normalization and clustering now have to be implemented considering all samples (see Supplement), from which diversity scoring can inform a ranking of intratumor-heterogeneity across samples. Single dots in the tSNE plots represent single cells, which might either be marked according to their cluster classification or according to their sample of origin.

**Figure 2.**
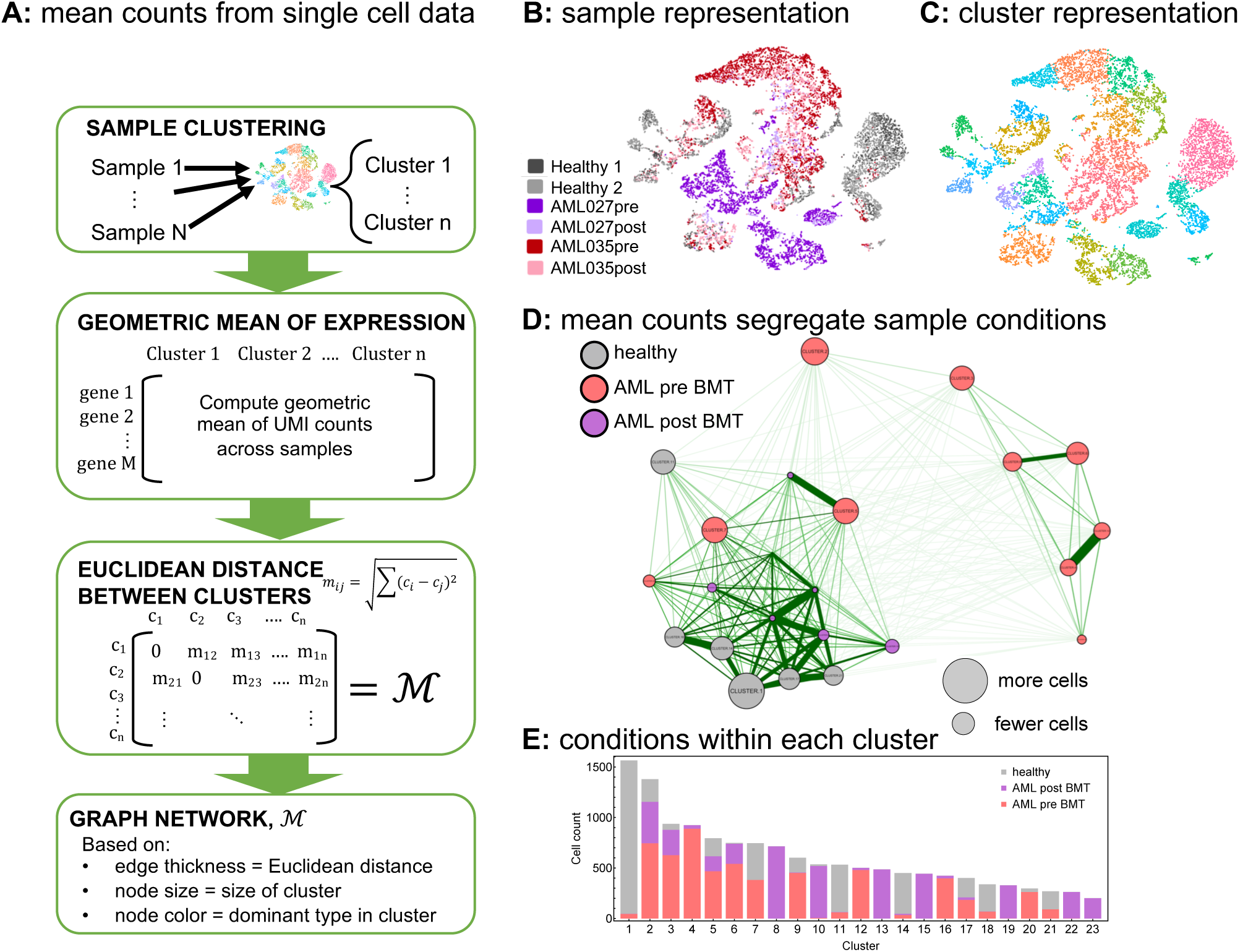
Mean cellular gene expression across clusters within patients can separate disease conditions to some degree. Here we built a network based on mean differences in overall expression. **A:** The geometric mean of UMI counts across samples and genes was calculated for each cluster. Then a Euclidean distance was calculated between clusters. Here we used publicly available scRNA-seq data^20^: two healthy donor BMMCs, two AML patient BMMCs pre bone marrow transplant (preBMT), and post bone marrow transplant (postBMT). These six samples were then clustered (using 10x-genomics Loupe™ Cell browser; for alternative clustering methods see Supplement), for which we show the sample-based **(B)** and cluster-based **(C)** tSNE-plots out of Loupe™ browser. Each dot represents a single cell, which is colored either according to sample of origin or its assigned cluster. The cluster-based differences in mean gene expression over unique molecular identifier (UMI) counts the gave rise to a “clustering of the clusters” **(D)**. The nodes in the resulting graph were colored based on the dominant cell type from each condition present in each cluster; gray for healthy, red for AML pre BMT and purple for AML post BMT, the distance between nodes was chosen inversely proportional to the difference in mean gene expression level, the individual distributions of cells from a specific condition in each cluster are shown in **E**.

Second, we sought to characterize across-sample differences by calculating the Kolmogorov-Smirnov (KS) distance^22^ of the cell count distributions in each cluster, in order to compare samples or pooled samples of the same condition (e.g. disease v. healthy) in terms of the cellular distribution over the identified clusters (**Fig. 3 A-G**).

**Figure 3.**
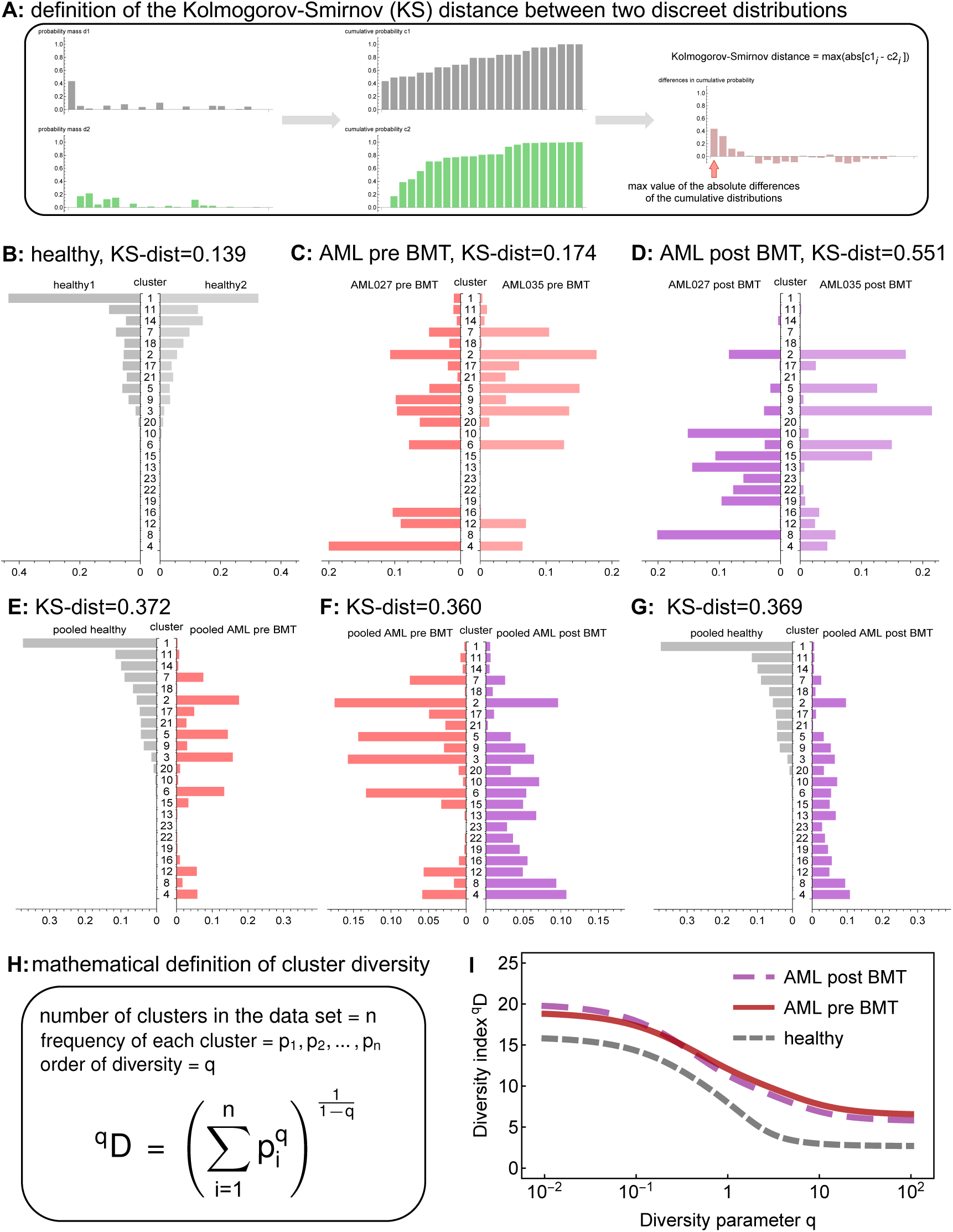
Cluster-based diversity scoring reveals strong differences between healthy individuals and cancer patients. In our analysis, using data from Zheng, et. al.^20^, we evaluated our ability to score significant differences in “cluster diversity” across healthy and AML samples pre and post bone marrow transplant (BMT). As a first indicator of between-sample or between-condition differences we used the Kolmogorov-Smirnov (KS) distance for discrete probability mass functions **(A)**. Little difference was found in the KS distance within condition-differences **(B, C)**, except in post BMT samples **(D)**. Between condition-differences were larger when comparing pooled samples across conditions **(E, F, G)**. We calculated a general diversity index (GDI) qD (H) to quantify diversity across “orders of diversity” q. For all orders of diversity measure, AML patients (pre and post BMT) had a higher diversity index compared to healthy individuals (**I**, 2 samples per condition), suggesting GDI can be used as a metric for stratification.

Third, we calculated an ecological diversity index^23^ using the cellular frequencies over clusters, across a range of order of diversity (**Fig. 3H, I**). To assess the robustness of our diversity metric, we performed down sampling of the original datasets and found the relative change in diversity index, across a range of order of diversity, to determine the sensitivity of our diversity metric (**Fig. 4**).

**Figure 4.**
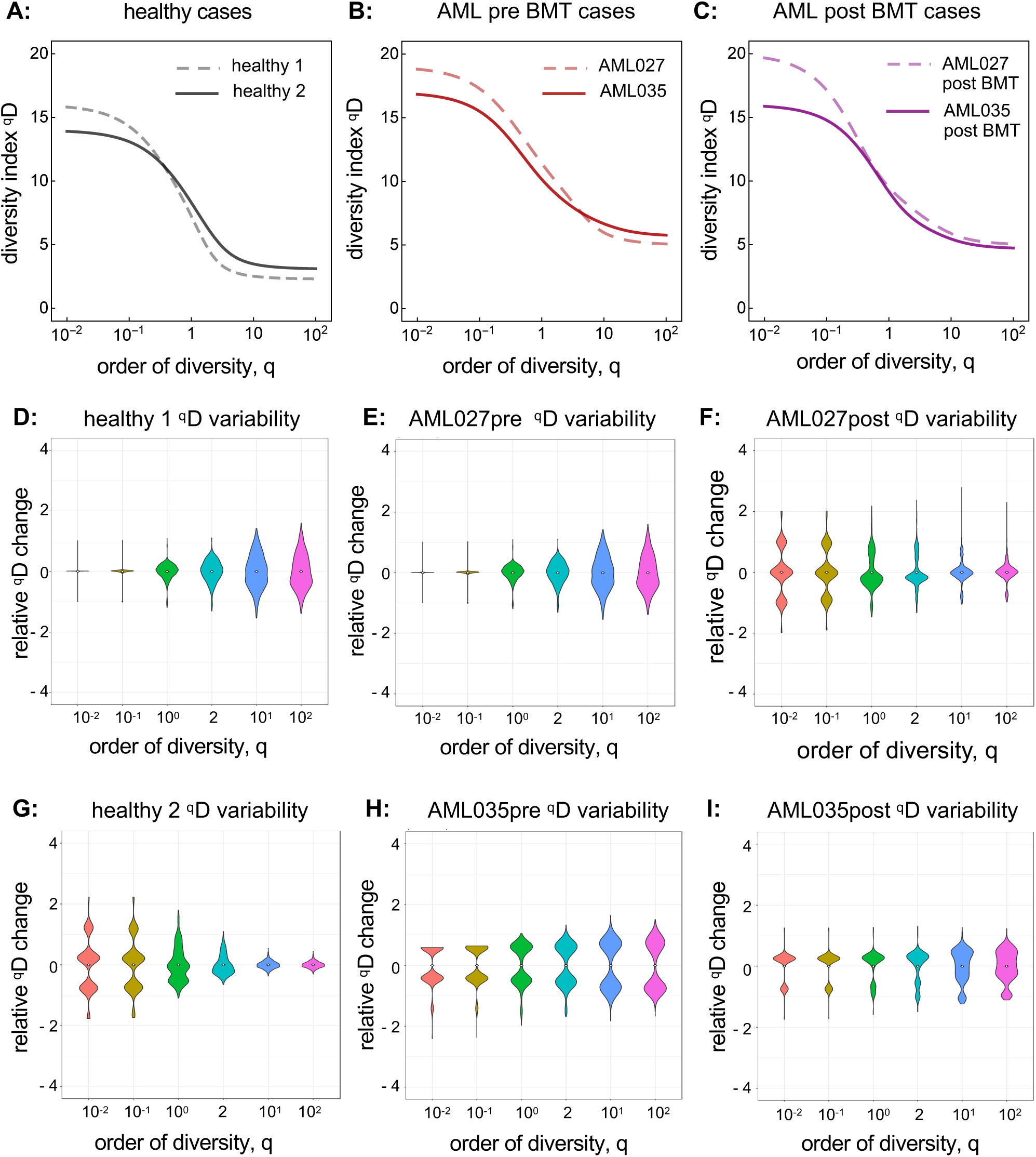
AML patients have consistently higher diversity compared to healthy individuals. Individual diversity spectrums were reported for healthy **(A)**, AML pre BMT **(B)**, and AML post BMT **(C)** samples (each line is one from one sample). Cell-gene matrices were down sampled to 50% of the cells 1000 times, and qD scores were calculated (using the full pipeline, see Supplement) for specific values of q=0.01, 0.1, 1, 2, 10, 100. The distributions of relative qD changes for healthy **(D, G)**, AML **(E, H)**, and postBMT AML **(F, I)** samples showed that generally, lower q values lead to less change in measured diversity. Across all cases the diversity score did not change by more than two units (relative change is measured by dividing the entire distribution by the distribution mean). For sample sizes, see Supplement. BMT: Bone marrow transplant.

Last, we applied our ITH pipeline and diversity metric to two additional datasets, (1) a hematopoietic cell type datasets comparing CD34+ cells with CD4+, CD14+, and CD19+ cell populations^20^, and (2) a lung cancer dataset with tumor-normal matched tissue sites taken from six different lung cancer patients^21^ (**Fig. 5**, Supplement). Further specific details of our methods, such as cells per sample, are described in the Supplement and is available online (including all code used to generate our results): https://github.com/mcfefa/scRNAseq.

**Figure 5.**
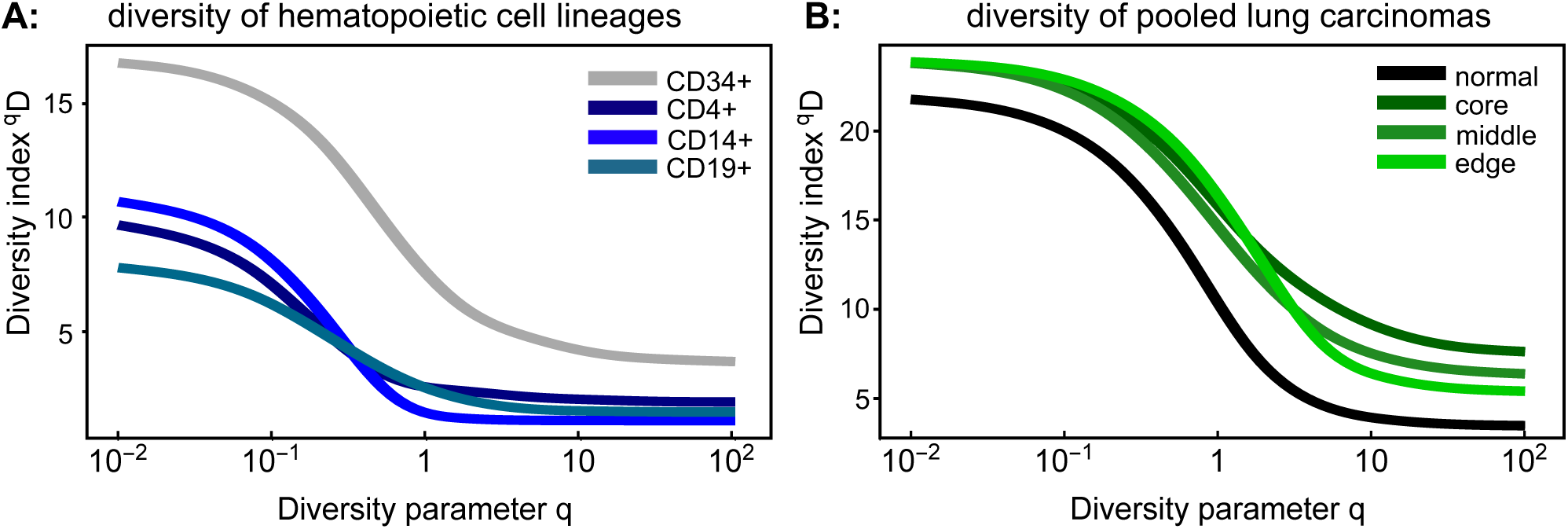
Higher diversity indicates higher clonality in in normal tissues and solid tumors. **(A)** Additional available data from Zheng, et. al.^20^ for CD34+ cells, CD4+ helper T-cells, CD14+ monocytes and CD19+ B-cells were run through pipeline S (see Supplement), and the continuum of diversity was calculated for each population. The naturally polyclonal population (CD34+) shows the highest diversity score. Each of the other differentiated immune cell compartments are more homogeneous across orders of diversity. **(B)** In solid tumors, location matters: normal-tumor matched lung carcinoma samples were obtained from publicly available data for six lung cancer patients^21^ (individual patients: see Supplement). The diversity metric across q shows an increase in diversity within tumors across different tumor locations.

## RESULTS AND DISCUSSION

We established proof of concept that we can generate clinically relevant summary metrics of ITH by analyzing publicly available scRNA-seq data^20,21^. Within BMMC samples from diagnosed AML and healthy control groups, we sought to establish how to summarize both inter- and intra-sample ITH. First, we clustered transcript expression of two healthy individuals, two AML patients, each sampled twice—once before and once after allogenic bone marrow transplant.

As a verification we sought to distinguish between healthy and AML samples based on the mean expression values across cells, across clusters (**Fig. 2A)**. With the 23 clusters identified (**Fig. 2B, C**), a network of clusters emerged, displayed as an undirected graph where the distance between mean UMI counts determines the thickness and length of the edges (**Fig. 2D**). The size of the node was chosen to indicate of the total number of cells in that cluster. We colored each node according to the condition (health, pre BMT AML, post BMT AML) that was in the majority (breakdown of actual proportions per cluster shown in **Fig. 2E**). This showed that indeed the large clusters with mostly healthy cells are most similar in average gene expression, while the large clusters with mostly AML cells cluster separately (to the right). The post BMT cells clustered more closely to the healthy than pre BMT AML samples. This supports the idea that these patients were potentially still transitioning but closer to a healthy phenotype. However, some AML-dominant clusters still grouped near the healthy/post BMT super cluster. Based on this “bulk” measure alone, one may not be able to easily distinguish between healthy and diseased cells. Therefore, other quantifications and metrics to describe the gene expression differences may better discriminate clinical and stages.

To determine metrics better at discriminating between healthy and disease AML, we analyzed and summarized inter- and intra-heterogeneity in two different ways. First, we considered each samples’ grouping of cells into clusters of similar gene expression. To this end, we used the Kolmogorov-Smirnov (KS) distance, which compares two discrete probability mass functions (the fraction of cells per cluster, **Fig. 3A**). We identified rather similar distributions within the same condition, and rather different distributions between conditions, with post BMT being a notable exception (**Fig. 3B-D**). The KS distance between the two healthy samples was 0.139, between the two pre BMT AML samples it was 0.174, but between the two post BMT samples it was 0.551. Since we had clustered all six samples together, we could also compare them pooled by condition, which revealed that indeed conditions distribute differently across the identified cellular subpopulations in high-dimensional gene expression space (**Fig. 3E-G**).

Second, we calculated a general diversity index (GDI) for each condition (healthy, pre BMT AML, post BMT). The mathematical definition of GDI, ^q^D, is shown in **Fig. 3H**. We established segregation of the different clinical conditions according to this ecology-based diversity index^23^. The pre BMT AML samples had consistently higher diversity index compared to the healthy sample and this held true across the entire order of diversity range, q (**Fig. 3I**). Interestingly, on this level the post BMT samples also scored unanimously higher in GDI. This could indicate that post BMT settings may require a certain amount of time after transplant to evolve toward a healthy spectrum of intra-leukemic diversity. Also, in a comparison of the individual samples within each condition (**Fig. 4, A-C**), the post BMT samples were most different from each other.

To interrogate the robustness of GDI further and to establish confidence in the metric, we down sampled the dataset and then re-clustered and calculated the ^q^D spectrum (**Fig. 4D-I**). During down sampling, we analyzed each sample individual by randomly removing 50% of the cells, then calculating the number of clusters identified for that individual’s transcript expression, and finally calculating the diversity index for specific q values of interest, including q=10^-2^, q=10^−1^, q=1 (which relates to the Shannon index), q=2 (which defines the inverse of the Simpson index), q=10, and q=10^2^. The distributions shown were obtained from 1000 runs of independent down sampling. Intriguingly, these distributions showed that with removing half of the cells, the diversity scores did not change more than a unit or two in either direction. When comparing to the diversity spectrum shown in **Fig. 3I**, this suggests that if healthy diversity spectrum were shifted up by two units (10% of the maximum) and the AML samples diversity spectrum were shifted down by two units, there would still be visible separation between the healthy and AML conditions.

To further validate our metric, we implemented on our approach with two other datasets. One dataset described different hematopoietic cellular subtypes, CD34+, CD4+, CD14+, and CD19+^20^. CD34+ is a hematopoietic progenitor cell marker and represent a polyclonal population that includes many different subtypes (hematopoietic stem cells, multipotent progenitor cells, common myeloid progenitor cells, common lymphoid progenitor cells, megakaryocytes erythroid progenitor cells and granulocytes macrophage progenitor cells) all of which express CD34^24–26^. The CD34+ polyclonal population contrasted the CD4+, CD14+, and CD19+ populations, which represent more homogenous cellular populations (helper T-cells, monocytes, and B-cells, respectively). This clonality pattern was recovered by GDI (**Fig. 5A**), where the CD34+ population had a considerable higher diversity score across the spectrum. Interestingly, lower values of q seem to separate differentiated cells or robustly.

Finally, we quantified ITH using lung cancer scRNA-seq samples^21^. We analyzed six different patients, each with up to three different tumor sites (core, middle, and edge) and a patient-matched adjacent normal lung tissue sample. Using our GDI metric, we see that the diversity spectrum of the normal lung tissue was much lower than any of the tumor site diversity scores (pooled conditions, **Fig. 5B)**. Interestingly here, more clear separation of conditions was achieved at high orders of diversity q, indicating differences in the number of driver clones at different sites within the tumors. These additional datasets further support the ability of quantified diversity metric to discriminated between healthy and diseases states, which can be applied in a clinical setting.

## SUMMARY AND CONCLUSIONS

Single-cell RNA sequencing efforts have greatly helped to uncover population structures and mapping to specific cellular population patterns^27^. Although these methods can also elucidate tumorigenesis^28^, immune-profiles^29^, and detect and track genomic profiles of clones^30,31^, the overall utility of scRNA-seq for cancer progression survival metrics has been elusive^32^. Here, we demonstrated the potential utility of two scRNA-seq based scores of cellular heterogeneities, using a generalized diversity index that may be elevated in disease. Remarkably, using previously published data without further processing, this quantification of intra-tumor heterogeneity was able to accurately distinguish AML from healthy individuals, as well as from post-transplant conditions. These data suggest that ITH can be estimated by diversity-based summary statistics, and that these summary statistics can be leveraged to predict clinical outcomes.

Our work here aimed at optimizing and identifying a clinically relevant summary index for ITH in the context of AML, for which targeted single-cell genome sequencing was also able to sensitively uncovered complex clonal evolution^33^. We anticipate that our ILH metric will be prognostic for leukemia-free survival (LFS) and potentially overall survival (OS), even after correction for known clinical prognostic variables. We have also shown how this metric can also be used to effectively describe heterogeneity in other malignancies, including solid tumors such as lung carcinomas.

From a clinical perspective in terms of tumor heterogeneity and emergence of resistance clones during targeted therapy^34,35^, we expect our metric can discriminate patients clinically. We hypothesize that these heterogeneity metrics would be elevated *independently* (at least *a priori)* in potentially highly resistant patients. The advantage of a more general metric used here is that it allows us to look across many orders of diversity, and potentially pick a desired range of heterogeneity quantification. For example, one might be interested especially in lower values of q, where a higher diversity score may indicate an individual (sample) more at risk of resistance evolution, as it shows high standing variation. On the other hand, differences at high values of q point to key differences in the number of important driver clones, which might uncover distinct vulnerabilities that can be targeted in combination or adaptively.

Diversity measures have long received attention in ecology and evolution^19,36^. We here approached to measure diversity, and thus tumor heterogeneity, using a general definition of non-spatial diversity^37^, in form of the quantity ^q^D (**Fig. 3H-I**). This approach considers all possible orders of diversity q, but also allows to compare disease stages according to a specific diversity index (fixed choice of q), which emerge as special cases of ^q^D. The species (clonal) richness of a sample is given by q=0. The Shannon index (log scale) can be found when q approaches 1. The Simpson index, which approximated the probability that any two cells are identical emerges from the case q=2. Both Shannon and Simpson index have been used in mathematical and statistical models of cancer evolutionary dynamics to quantify tumor heterogeneity as it potentially changes during tumor growth, with disease progression, or during treatment^38–40^. Shannon entropy-based statistics have also been used to quantify single cell heterogeneity, to deliver insights into emerging or disappearing clones during transitions between clinical conditions^41^.

Single cell RNA sequencing experiments give a snapshot of the cell population state on the level of gene expression, and it can characterize how individual cell’s transcriptomes compare to the bulk. In contrast to mass cytometry, drop-seq is fast and extremely high-throughput. Other single cell technologies, like flow cytometry can be used to generate single cell data for a relatively small subset of potential markers that distinguish between normal and disease, and it requires that the researcher/clinician knows what these markers are in advance. Among a variety of outcomes that may be distinguished by our metric, one can be the study for segregating samples based on disease severity, for which additional follow-up knowledge will be needed.

Further extending our results to the potential impact in a clinical setting in leukemias, intraleukemic heterogeneity (ILH) is a known reservoir for tumor resistance and clinical refractoriness to targeted therapies^9^. For current targeted therapies to treat AML, clinical responses have been modest. Specific means to change ILH can be of particular appeal in such cases, as they might help render tumors less aggressive and hinder their ability to rapidly evolve resistance. In particular, hypomethylating agents (HMA, cytosine analogs that irreversibly bind to DNA methyltransferase, an enzyme required for methylation of CpG-rich DNA) have the potential to diminish ILH^42^. Transcriptome changes upon treatment with HMA therapy have not been analyzed at single cell resolution. Our analyses here provide a quantitative basis to understand and reliably track these changes.

Clear separation of diversity metrics by condition, as we show it, might not be expected in general. A weakness of our approach is that it does not consider any meaning of the associated phenotypes or genotypes. Therefore, as it stands, our method cannot be transferred to improve the predictive power of existing bulk signatures. Hence, existing survival data is unlikely to be useful to prove that GDI is predictive of survival, and novel databased that uniquely connect high throughput single cell experiments with clinical outcomes are needed.

Once the appropriate cohorts are established, however, changes in an individual’s diversity score could indicate unique features of disease progression. In the context of adaptive therapy^43,44^, which aims at tumor burden control rather than difficult tumor eradication, it might be critical to identify the appropriate scale of diversity that best predicts outcomes. One could speculate that there is an optimal window of diversity that should be maintained—very low diversity could indicate fast disease progression and very high diversity could mean that the tumor could adapt to the treatment schedule too quickly. The concept we introduced here is sufficiently flexible in its ability to quantify optimally predictive windows of diversity that should be maintained during adaptive therapy.

## AUTHORSHIP CONTRIBUTIONS

P.M.A., E.P. conceived the study. M.C.F., M.B. and P.M.A. performed computational experiments and statistical analyses. M.C.F., and P.M.A. performed mathematical modeling. M.C.F., M.B., E.P. and P.M.A. analyzed the data. M.C.F., M.B., E.P. and P.M.A. wrote the manuscript. P.M.A. and E.P. supervised the project.

## SUPPLEMENTAL MATERIAL

### SUPPLEMENTAL MATERIALS AND METHODS

#### DROPLET-BASED scRNA_SEQ_ SAMPLES

The 10X Genomics platform offered a variety of datasets^1^ that were used to create our pipeline for quantifying intraleukemic heterogeneity. The major focus of our analysis and pipeline development was of Healthy/Control and AML patient bone marrow mononulear cells (BMMCs). The Healthy Controls 1 and 2 BMMCs had been sequenced on Illumina Hiseq 2500 Rapid Run V2 with 90–135 thousand reads per cell and 2000–2400 cells detected. The AML027 and AML035 pre-transplant BMMCs had been sequenced on Illumnia NextSeq 500 High Output with 16.6–58 thousand reads per cell and 3500–3900 cells detected. The AML027 and AML035 post-transplant BMMCs had also been sequenced on Illumina NextSeq 500 High Output with 41–51 thousand reads per cell and 900–3900 cells detected. Additional datasets^1^ for CD34+, CD14+, CD19+, and CD4+ (hematopoietic cell type set) cells as well as patient normal-tumor matched lung cancer samples^2^ available in ArrayExpress under accessions E-MTAB-6653 and E-MTAB-6149 were used to test the robustness of the pipeline. The CD34+ dataset was CD34+ cells enriched from peripheral blood mononuclear cells (PBMCs), sequenced on an Illumnia NextSeq 500 High Output with 24.7 thousand reads per cell with 9000 cells detected. The CD14+ dataset was enriched from PBMCs, sequenced on Illumnia NextSeq 500 Output with 100 thousand reads per cell with 2600 cells detected. The CD19+ dataset was enriched from PBMCs, sequenced on Illumnia NextSeq 500 Output with 25 thousand reads per cell with 10000 cells detected. The CD4+ dataset was enriched from PBMCs, sequenced on Illumnia NextSeq 500 High Output with 21 thousand reads per cell with 11000 cells detected. All lung tissue samples were prepared as single-cell suspensions, sequenced on Illumina HiSeq4000, and aimed for an estimated 4000 cells per library^2^. There were six patients analyzed, each patient had 4 distinct tissues locations sampled: an adjuacent normal lung sample (normal), a sample from the tumor core (core), a sample from the tumor margin (edge), and a sample between the tumor core and margin/edge (middle).

#### INITIAL DATA PROCESSING

First, the all samples were run through the 10x Genomics cellranger count pipeline for transcriptome alignment, calling individual cell barcodes, estimating multiplet rates, and finally clustering of normalization corrected FPKM (fragments per kilobase million; expression level) values (**Fig. S1**, pipeline X). In our analysis of the Healthy versus AML high-throughput scRNA-seq, we used cellranger aggr to pool together the individual dataset results, so we could compare intra-tumor heterogeneity (ITH) between samples with the same clustering. Using a graph-based clustering algorithm, 23 different clusters were identified based on clustering cells by expression similarity (shown in **Fig. 2B-C**). The clustering was performed by forming a graph with cells as vertices and edges indicating pairs of cells that are sufficiently similar (implemented by CellRanger). The CellRanger graph-based clustering with Louvain Modularity Optimization^3^ was used to partition this graph into clusters of similar cells. For analysis with the hematopoietic cell set and the lung cancer samples the cell/gene matrices produced by cellranger count pipeline were then loaded into R using the Seurat package^4^ (designed for QC, analysis, and exploration of single cell RNA-seq data; **Fig. S1**, pipelines M and S). With Seurat, cells were removed that had either fewer than 200 UMIs and greater than 6000 UMIs and more than 10% mitochondrial DNA^2^, then scaled and normalized before clustering using graph-based clustering.

#### QUANTIFYING SUMMARY DIVERSITY METRICS

After the data was clustered, we then sought to quantify whether the clustering could disentangle healthy and AML samples by average gene expression. To approach within-sample differences in overall gene expression, we computed Euclidean distance matrices of the mean expression values in each cluster, to establish a distance metric of cluster differences and a subsequent diversity of distances across samples (standard deviation, ANOVA) (**Fig. 2A**). For each cluster, the geometric mean of each unique molecular identifier (UMI) across all genes were computed, then the Euclidean distance was computed between clusters. This was plotted as a graph where each node represents each clusters identified in the leukemic dataset (**Fig. 2D**). The size of the node indicates the total number of cells in that cluster and the color identifies the major species (AML, Healthy, or postBMT) present in that cluster. The distribution of cells per cluster (**Fig. 2E**) was also included to show which other conditions were also present in each cluster and how the number of cells compared to the major species used to dictate the color of the nodes in graph describing the connectivity of the clusters in **Fig. 2D**.

Next, we sought to characterize across-sample differences by calculating the Kolmogorov-Smirnov (KS) distance^5^ of the cell count distributions in each cluster (**Fig. 3A**). The KS distance is a non-parametric measure that is calculated as the supremum of paired differences of two empirical probability mass functions^5^. Last, we calculated a continuum of the ecological diversity index^6^ based on the individual cell frequencies in each of the clusters identified by the graph-based clustering (**Fig. 3H**), across all clusters, which can be written as:

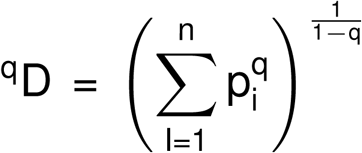
where n is the number of clusters in the data set, p_n_ is the frequency of each cluster, and q is the order of diversity. q is a hyperparameter that would be optimized in a clinical setting. The two distributions for AML were joined together and the two distributions of the healthy indiviuals were joined together and plotted across orders of diversity (**Fig. 4A-C**). The same technique was used to group the lung cancer samples based on location as reported in (**Fig. 5B**).

#### DOWNSAMPLING

We sought to test the robustness of our metrics by downsampling the number of cells and re-calculating the number of clusters identified as well as changes in key diversity scores. This downsampling was preformed removing increments of 10% of the data at a time and the results were reported as the summary statistics of 1000 runs at each downsampling percentile (**Fig. S2**). The downsampling results were also used to quantify confidence in the diversity index spectrum and was calculated for the following orders of diversity: 10^−2^, 10^−1^, 10^0^, 2, 10^1^, 10^2^. The relative change in diversity score was reported for the downsampling results with 50% the initial amount of cells in **Fig. 4D-I**. Relative change in the diversity score was calculated by subtracting the mean of the diversity score from all 1000 and reported as the distributions of scores around that mean score.

#### CODE AVAILABILITY

The code used in the pipelines described in **Fig. S1** were uploaded to a GitHub repository (https://github.com/mcfefa/scRNAseq). These pipelines were implemented using data run through at least cellranger counts, and then post-processed with CellRanger Loupe Browser, Mathematica, and R. Code available includes:

- Mathematica code and R scripts and data described to implement all pipelines (described in **Fig. S1**)
- Mathematica code for calculating the geometric mean of UMIs and Euclidean distances between clusters (for **Fig. 2**)
- R code to draw the graph of the clusters (**Fig. 2D**)
- Mathematica code for calculating the KS distance and diversity spectrum across all clusters for the leukemic dataset (**Fig. 3**)
- R code used to downsample datasets, then cluster and calculate diversity indices (**Fig. 4** and **S2**)
- R code for calculating the diversity spectrum for the hematopioteic subtype and matched lung carcinoma datasets (**Fig. 5**)

**Supplemental Figure 1.**
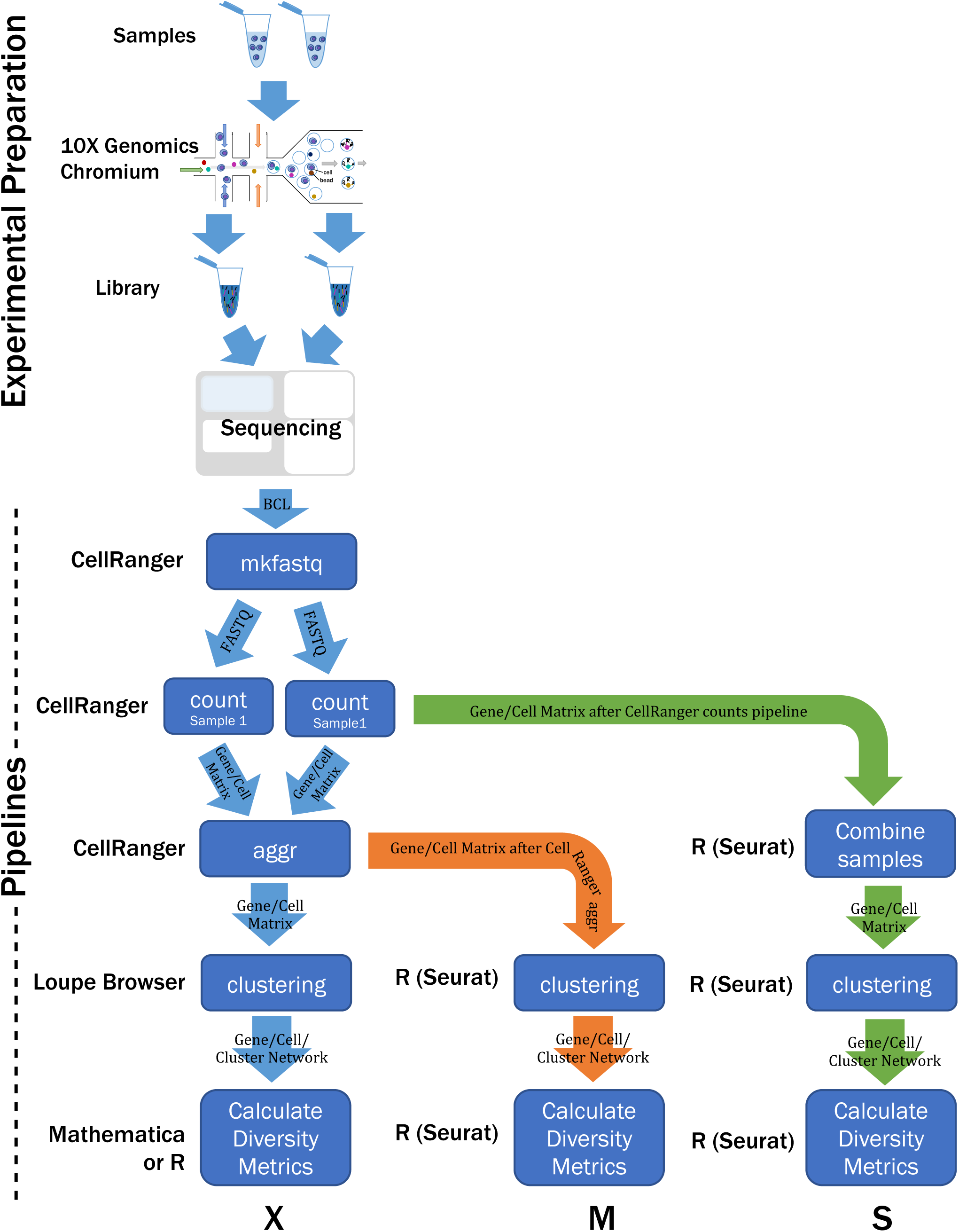
Workflow diagram for the scRNAseq quantifying intraleukemia heterogeneity for an example comparing two samples. Cells were processed through the 10X Genomics Chromium and ultimately sequenced, producing a raw base call (BCL) file, which was demultiplexed for each flowcell directory and converted to a FASTQ file using cellranger mkfastq. Then cellranger count was run separately for each library to align, filter, and count barcodes and UMIs. These instances were then aggregated into a single instance using cellranger aggr to normalize runs to the same sequencing depth and recompute analysis on combined dataset. Following along pipeline X, from the aggregated analysis, the Loupe Browser contains the clustered data, which were exported and run through a Mathematic script to calculate the diversity metrics. This pipeline was used to generate Figs. 3 and 4. This pipeline was expanded to give the user increased flexibility by using the gene-barcode matrices output by cellranger and then processing the data in R using the Seurat package, which allows users to set the filtering criteria (M) and allows data to be combined without running through the cellranger aggr pipeline (S) as well as both new pipelines allow users to implement additional clustering algorithms. Pipeline S was used to generate Fig. 5 from gene-barcode matrices available from additional data from Zheng, et. al.^18^.

**Supplemental Figure 2.**
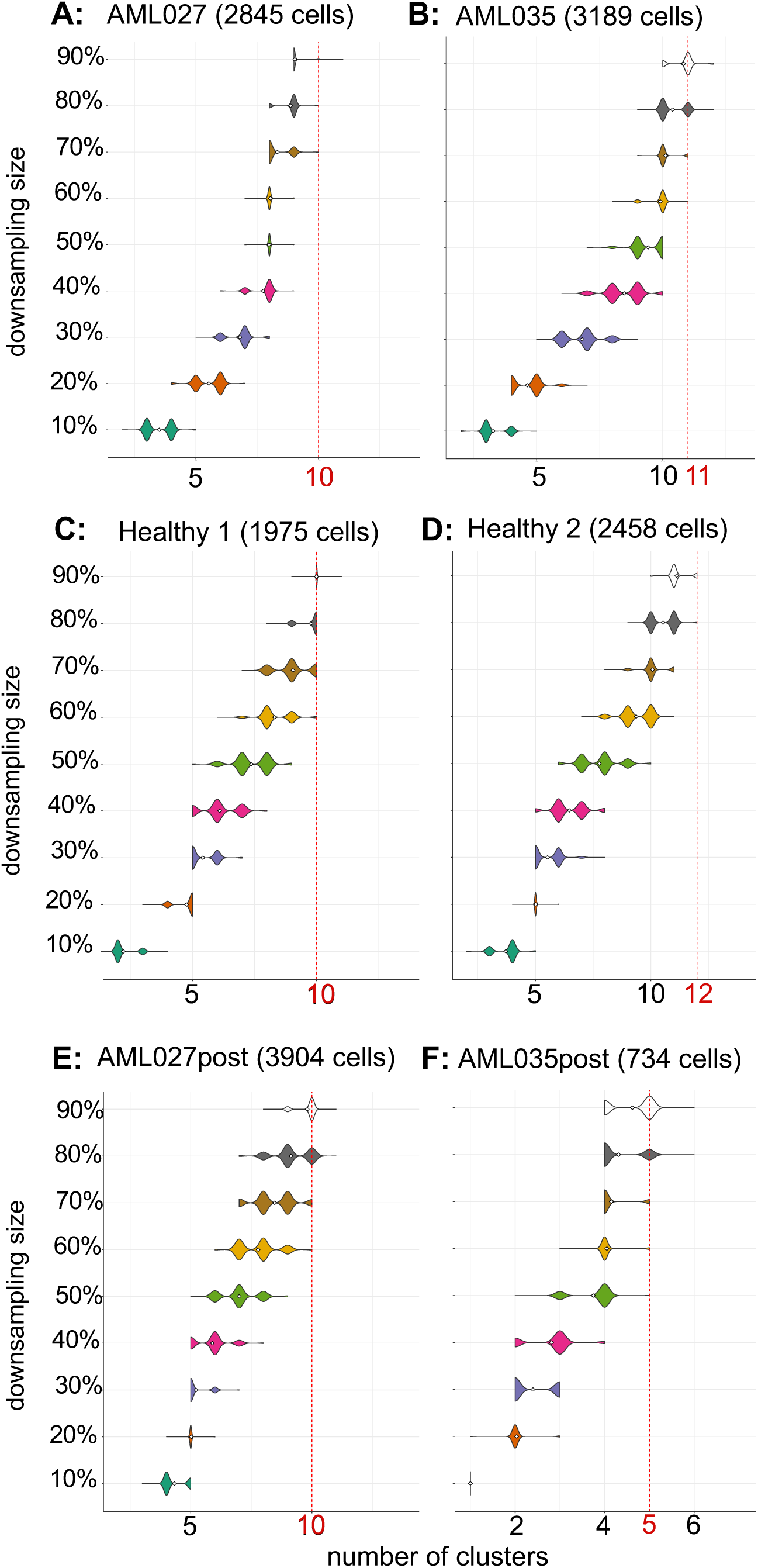
AML populations converge to consistent number of clusters that describe the population sooner than healthy populations. graph-based clustering. Each original population’s gene-barcode matrix obtain from publicly available data from Zheng, et. al.^18^ was then downsampled to contain 10% to 90% of the cells originally in the matrix. This data was loaded into R using the Seurat package, downsampled, and then clustered to determine how sensitive the cluster metrics was to the starting number of cells. Downsampling was performed 1000 times per cutoff and the violin plots show the distribution of clusters identified. The red-dashed lined indicates the number of clusters identified with no cells removed from the dataset. Gray numbers indicate the number of cells present in each downsampling condition. AML populations (**A**, **B**) showed quickest convergence to number of clusters after about 1500 cells were present in clustering. Healthy populations (**C**, **D**) showed that as more cells were added, an additional cluster could be found. AML post-bone marrow transplant populations (**E**, **F**) behaved more similar to healthy than AML populations.

**Supplemental Figure 3.**
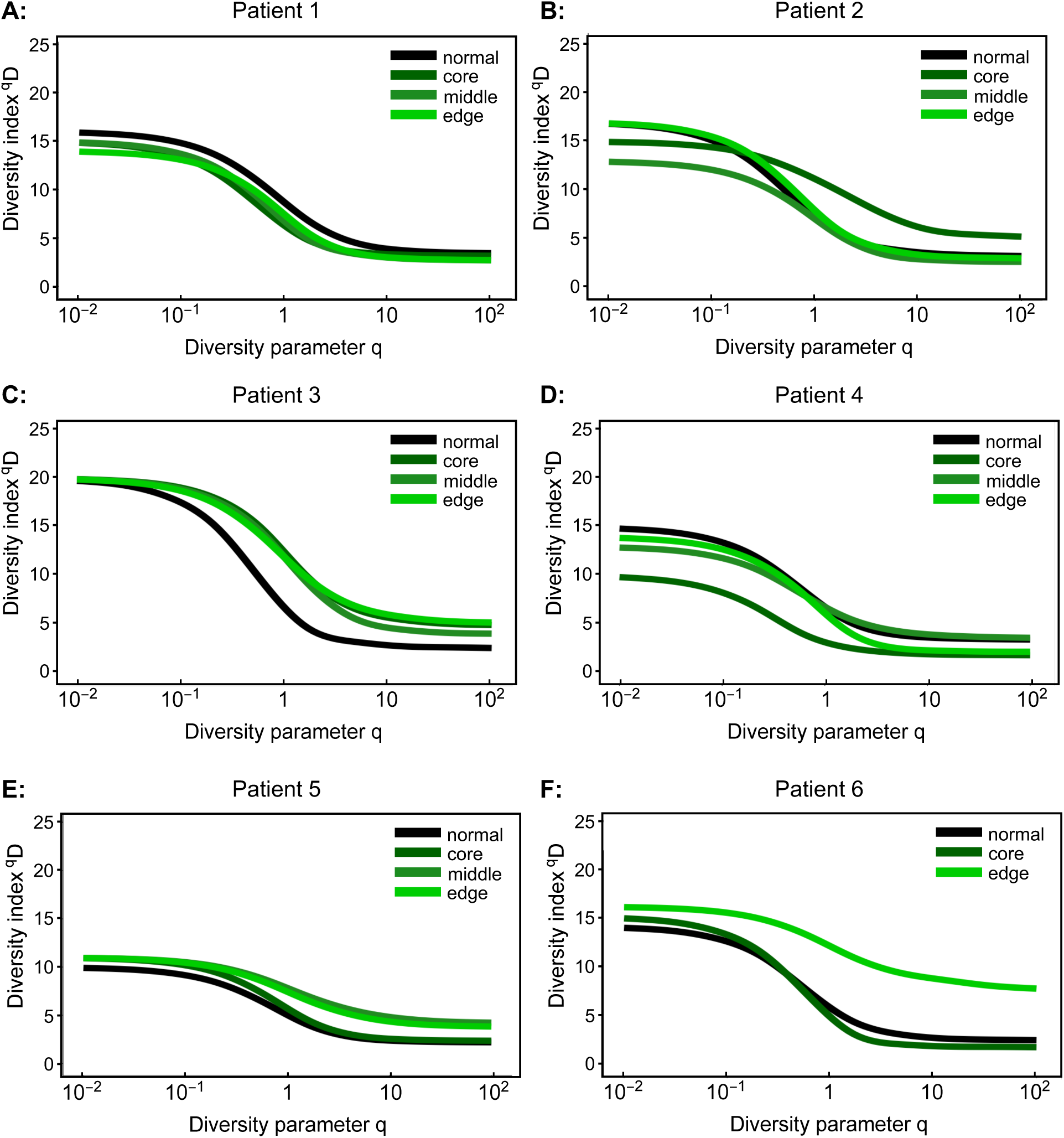
Individual lung cancer patient diversity scores show same trends as aggregated clustering results with tumor samples have greater diversity scores than their matched normal counterpart. Individual patients were run through pipeline S to aggregate normal, core, middle, and edge tissue samples.

